# Sex, not familiarity, shapes social interactions in adult captive tortoises (*Testudo hermanni*)

**DOI:** 10.1101/2025.07.23.666358

**Authors:** Silvia Damini, Elisabetta Versace, Gionata Stancher

**Affiliations:** Center for Mind/Brain Sciences, University of Trento, Rovereto, Italy; Department of Behavioural and Cognitive Biology, University of Vienna, Vienna, Austria; School of Biological and Behavioural Sciences, Centre for Brain and Behaviour, Queen Mary University of London, London, UK; Rovereto Civic Museum Foundation, Rovereto, Italy

**Keywords:** solitary species, social cohesion, social behaviour, recognition, sex differences, tortoises, *Testudo hermanni*

## Abstract

Social cohesion varies across species. Solitary animals minimise social contact to reduce costs such as resource competition, aggression, and disease transmission. Solitary *Testudo* hatchlings can rely on familiarity to avoid unfamiliar conspecifics. It remains unclear whether this strategy is used in adult captive tortoises and whether sex differences exist. To address these questions, we experimentally investigated a colony of captive *Testudo hermanni* tortoises (a vulnerable species), housed in outdoor enclosures. After a group familiarisation phase, we tested adult tortoises in a novel arena with either a familiar or an unfamiliar same-sex individual (not encountered for at least 3 months). Contrary to our expectations, familiarity had no significant effect on social cohesion measures such as inter-individual distance or facing orientation, showing that in adult captive tortoises familiarity had limited effect on social cohesion. In contrast, we found striking sex differences: males approached conspecifics quickly and remained in close proximity throughout the test, frequently displaying aggression; females initially explored the partner but then distanced themselves, with minimal aggression, similarly to what *Testudo* hatchlings have been observed to do with unfamiliar individuals. These results show that in adult captive tortoises adult social interactions and social cohesion are influenced by sex. This study provides insight into social cohesion and interaction strategies in tortoises, relevant for captive settings and conservation planning.

## Introduction

Social cohesion is a fundamental life-history trait that shapes individual fitness and survival via species-specific tendencies to maintain proximity and coordinate within a group (Krause and Ruxton 2002; Silk 2007; Salguero-Gómez 2024). Species differ widely in their social cohesion, ranging from social taxa that live in close proximity and form stable, organized groups, to solitary species that restrict social interactions to specific contexts such as mating or territory defence (Hofmann et al. 2014; Rubenstein and Abbot 2017; Twining et al. 2024; Makuya and Schradin 2024; Salguero-Gómez 2024). While social cohesion may benefit animals in terms of more efficient antipredator strategies, thermoregulation, social learning (Ward and Webster 2016), or reduced energetic costs (Ebensperger and Cofré 2001), it also imposes costs such as competition for limited resources, increased risk of disease transmission, and aggression (Krause and Ruxton 2002). In solitary or non-gregarious species, minimizing social contact may represent an adaptive strategy in response to ecological pressures and energetic trade-offs, such as wide distribution of food sources and predator avoidance based on camouflage (Makuya and Schradin 2024). Depending on the context, the reduction of costly encounters may rely on cognitive mechanisms such as general conspecific avoidance, the ability to distinguish between familiar and unfamiliar conspecifics or individual recognition (Tibbetts and Dale 2007). Individual recognition entails the ability to identify a specific individual, as opposed to familiarity recognition, where is sufficient to recognize a more general group specific label (Ward et al. 2020).

Recognition of familiar individuals has been demonstrated across diverse taxa, including insects (Tibbetts and Dale 2007; Sheehan and Tibbetts 2011), birds (Versace et al. 2021), mammals (Cheney and Seyfarth 1980; Kendrick et al. 2001), amphibians and reptiles (Versace et al. 2018; Munch et al. 2018; Ray and Maruska 2023), in the context of both affiliative and avoidance/agonistic strategies (Husak and Fox 2003; Versace et al. 2018). A well-documented example is the “dear enemy effect”, where animals display lower levels of aggression towards known conspecifics (usually neighbours) compared to unfamiliar ones, with the advantage of saving energy of reaffirming already determined social hierarchies and functioning behavioural modalities (Ydenberg et al. 1988). This phenomenon has been observed in various taxa, including territorial amphibia (Bee and Gerhardt 2002) and reptiles (Aragón et al. 2003; Versace et al. 2018).

Despite historical underrepresentation in the study of social behaviour and cognition, reptiles are increasingly recognized for their social competencies (Doody et al. 2013, 2021), including non-random social networks, with the discovery of interaction opportunities, for instance in the sharing of burrows in tortoises (Sah et al. 2016; Madile Hjelt et al. 2024). Tortoises represent a compelling model for the study of social behaviour in ectotherm species due to their largely solitary lifestyles, precocial development, and for being one of the most threatened vertebrates worldwide (Rhodin et al. 2018; Chen et al. 2025). As social behaviour has been connected to the spread of disease in tortoises (Wendland et al. 2010), and tortoises are often housed with conspecifics in zoos and conservation facilities (Freeland et al. 2020; O’Brien et al. 2025; Goessling et al. 2026), ecological and practical implications are associated with the study of social behaviour in tortoises.

*Testudo hermanni*, the species investigated in our study, is a species of concern listed as “vulnerable”(IUCN 2024). *T. hermanni* is not known to form pair bonds or cohesive groups and shows limited interactions restricted to reproductive contexts such as courtship, mounting, and nesting (Auffenberg 1977; Ernst and Barbour 1989; Pearse and Avise 2001; Sacchi et al. 2003; Galeotti et al. 2005). During courtship, male tortoises of this species use physical and vocal courting signals that indicate individual quality and influence female choices (Galeotti et al., 2005). As many other reptile species (Mason & Parker, 2010), Hermann’s tortoises also communicate through chemosensory and olfactory cues, with both sexes being able to discriminate between their species and other species’ odours, while only males can use olfactory information to infer sex and sexual maturity of conspecifics, showing that males and females might rely on different communication cues (Galeotti et al., 2007). This indicated sexual dimorphism in olfactory sensitivity, suggesting that in males and females might rely on different communication channels during social interactions. Sex differences have been observed also in home range (Mazzotti et al. 2002). Much less is known on social cohesion differences between sexes.

Natural population densities of *T. hermanni* rarely exceed 10 individuals per hectare (Stubbs and Swingland 1985). Yet, in conservation sanctuaries and other captivity settings, tortoises are frequently housed at much higher densities, in conditions that prompt more frequent interactions with conspecifics (Burghardt 2013; Freeland et al. 2020). Under such conditions, captive tortoises have more chances of interacting with conspecifics than wild ones, raising questions on social interaction and recognition of familiar individuals in captive contexts. Research on captive tortoise has revealed that adult tortoises housed together with conspecifics can follow the gaze of conspecifics (Wilkinson et al. 2010b) and learn from their actions (Wilkinson et al. 2010a). Studies in controlled settings have revealed social competencies in tortoises. Familiarity-based discrimination has been observed in juveniles of closely related species (*Testudo graeca* and *T. marginata*), which ignore familiar individuals, but actively avoid unfamiliar individuals after a first exploration (Versace et al. 2018). This strategy can facilitate dispersal of tortoise clutches, diminishing competition for resources (see also Goessling et al. (2026)). We are not aware of studies on familiarity-based discrimination conducted in *T. hermanni*. Moreover, it remains unclear whether such social avoidance strategies are used by adults in captive settings. Tracking of some Chaco tortoises (*Chelonoidis chilensis*) (Hjel at al., 2024) and desert tortoises (*Gopherus agassizii*) (Sah et al. 2016) that highlight shared space use and non-random social networks, call for experimental studies to better understand social interactions and recognition in tortoises.

Regarding sex differences in reptiles, in the amphisbaenian *Trogonophis wiegmanni*, females but not males discriminate between familiar and unfamiliar juveniles (showing a higher tongue flicker rate) using chemosensory cues (Martín et al. 2021). Sex differences have also been found in *T. hermanni* in habitat use (Mazzotti et al., 2002), and in inhibitor control, with females outperforming males in laboratory settings (Santacà et al. 2025). It remains to be clarified whether sex differences exist in encounters with conspecifics, and what dynamics emerge in captive settings.

In the study presented here, we used captive adult *T. hermanni* raised outdoors in a semi-natural station to understand whether these animals discriminate between familiar and unfamiliar same-sex conspecifics, and whether this discrimination affects social cohesion/distancing and aggression. Based on previous results on tortoise hatchlings of the closely relate species *T. graeca* and *T. marginata* (Versace et al. 2018), we hypothesised that familiar and unfamiliar pairs might behave differently, with familiar pairs ignoring the other individual, and unfamiliar individuals actively moving away from one another after a first exploration, as observed in hatchlings. In our previous study we could not determine the sex of the hatchlings, for no sexual dimorphism is present at such young age. Because in this study we expanded to adults, we could investigate the role of sex in shaping social cohesion and social interactions between familiar and unfamiliar tortoises. As studies in the wild often suffer from high environmental variability, our semi-controlled setting offered an opportunity to expand knowledge on tortoises, and inform conservation efforts.

## Methods

The experiment consisted of two phases: (1) a familiarisation phase, during which tortoises were housed with same-sex conspecifics in an outdoor enclosure, and (2) a test phase conducted in a neutral novel arena indoor. During familiarisation, tortoises were kept in groups of four individuals of the same sex for a minimum of two months to allow social recognition to develop. Each subject was subsequently tested in two trials: once with a familiar individual (familiar condition) and once with an unfamiliar individual (unfamiliar condition). The order of conditions was counterbalanced across subjects (see Table 1 for a full test schedule). During each trial, we recorded the inter-individual distance, relative orientation, and instances of aggressive behaviour (data available in Table 5).

**Table 1.** List of experimental Pairs, their Sex, and Familiarity condition they were tested in.

| Familiarity condition | Sex | Pair |
| --- | --- | --- |
| Familiar | Male | 2-9 |
| Familiar | Male | 6-18 |
| Familiar | Male | 11-12 |
| Familiar | Male | 1-3 |
| Familiar | Male | 7-8 |
| Familiar | Male | 4-5 |
| Unfamiliar | Male | 1-4 |
| Unfamiliar | Male | 2-8 |
| Unfamiliar | Male | 3-5 |
| Unfamiliar | Male | 7-9 |
| Unfamiliar | Male | 11-18 |
| Unfamiliar | Male | 6-12 |
| Familiar | Female | 35-36 |
| Familiar | Female | 25-32 |
| Familiar | Female | 26-30 |
| Familiar | Female | 33-34 |
| Unfamiliar | Female | 26-33 |
| Unfamiliar | Female | 30-34 |
| Unfamiliar | Female | 32-36 |
| Unfamiliar | Female | 25-35 |

### Subjects and rearing conditions

Experimental observations were conducted at the station SperimentArea, a semi-natural facility of the Rovereto Civic Museum (Rovereto, Italy) that works as a sanctuary for tortoises. We observed pairs from a total of 20 adult tortoises (*Testudo hermanni*): 8 females and 12 males, see Table 1. This relatively small sample size is noteworthy for a species listed as vulnerable (IUCN 2024). We assessed the size of tortoises measuring the plastron length from the central part of the gular scutes to the central part of the anal scutes (male size: average=15.8 cm, min-max=14.5-17 cm; female size: average=15.6 cm, min-max=12-18 cm). Tortoise estimated age, based on carapace size and degree of wear, ranged between 10 and 50 years old.

Tortoises are not endemic to this region. Animals recovered by the local forestry authorities were relocated to the sanctuary. Some animals were found as juveniles, some as adults. Tortoises were either domestic animals donated to the facilities, domestic animals escaped from captivity, or poached in the wild and then grown in captivity. Morphological characteristics observed in the subjects indicate a growth rate compatible with being raised in captivity (Klaphake 2010; Semaha et al. 2025). Animals had entered SperimentArea at least 3 years before the start of the experiment (typically 5 or more). We therefore consider this a captive tortoise population.

After the brumation period (November-end of March), and before the beginning of the experiment, animals were housed in same-sex groups in fenced areas of about 300 m^2^ each (15-20 m^2^ for each animal), with opportunities to explore, hide and dig burrows. At the beginning of the experiment (May), animals from the same fenced area were divided into five same-sex groups of four individuals, with random group allocation. Each group was housed for an average of 72 days (range: 60-89 days), with differences due to external weather, as animals were tested only in sunny days. This duration was considered sufficient to induce recognition of familiar individuals, as *Testudo* hatchlings succeeded in a similar task after only 3 weeks of exposure (Versace et al. 2018). Considering the brumation period and the familiarization period, unfamiliar individuals had not met each other for at least 3 months (tipically longer). As animals had entered SperimentArea as adults, and had previously rotated in space allocations, it was not possible to confirm the complete unfamiliarity of the subjects. However, the familiarity in the home enclosure was fully controlled. Males were housed in a 2.2×4.1 m lawn-like enclosure with a shelter; females were housed in a slightly larger 3×5 m enclosure containing a shelter and natural vegetation (e.g., bushes) to accommodate nesting behaviour. In these enclosures, all individuals were always visible to each other, while at the same time having enough space to move away from conspecifics in case of aggression. During the first days of group housing in the home enclosure, animals were observed to prevent/limit aggressive interactions. No human intervention or separation was required. Water was continuously available, fresh green leaves and vegetables were provided daily in one location, to maximize the chances of the tortoises meeting the other group members. Due to the limits of a relatively sample size, it has not been possible to run a full between-subjects design. All subjects have been tested twice: once in the familiar condition, once in the unfamiliar condition.

### Experimental apparatus

We tested tortoises in a dedicated room with no heating or air conditioning, in a plastic circular arena (diameter 102 cm; wall height 28 cm, see Figure 1) with the bottom covered in gravel. Excrements were cleaned out after each test. We placed a black plastic panel (height: 128 cm) around the arena to minimize distractions from the external environment. The apparatus was illuminated from above by a halogen light (500 W), suspended 205 cm from the ground. A Windows LifeCam camera (standard webcam with HD option) was secured above the centre of the arena to record behaviour. All tested were conducted indoor. Although the overall temperature fluctuated based on external weather, the halogen light kept it above 26°C, and tortoises comfortably moved in the arena. The subjects were not habituated to the arena before the trials. As they were tested only twice in the arenas, this was a relatively new environment during all tests.

**Figure 1.**
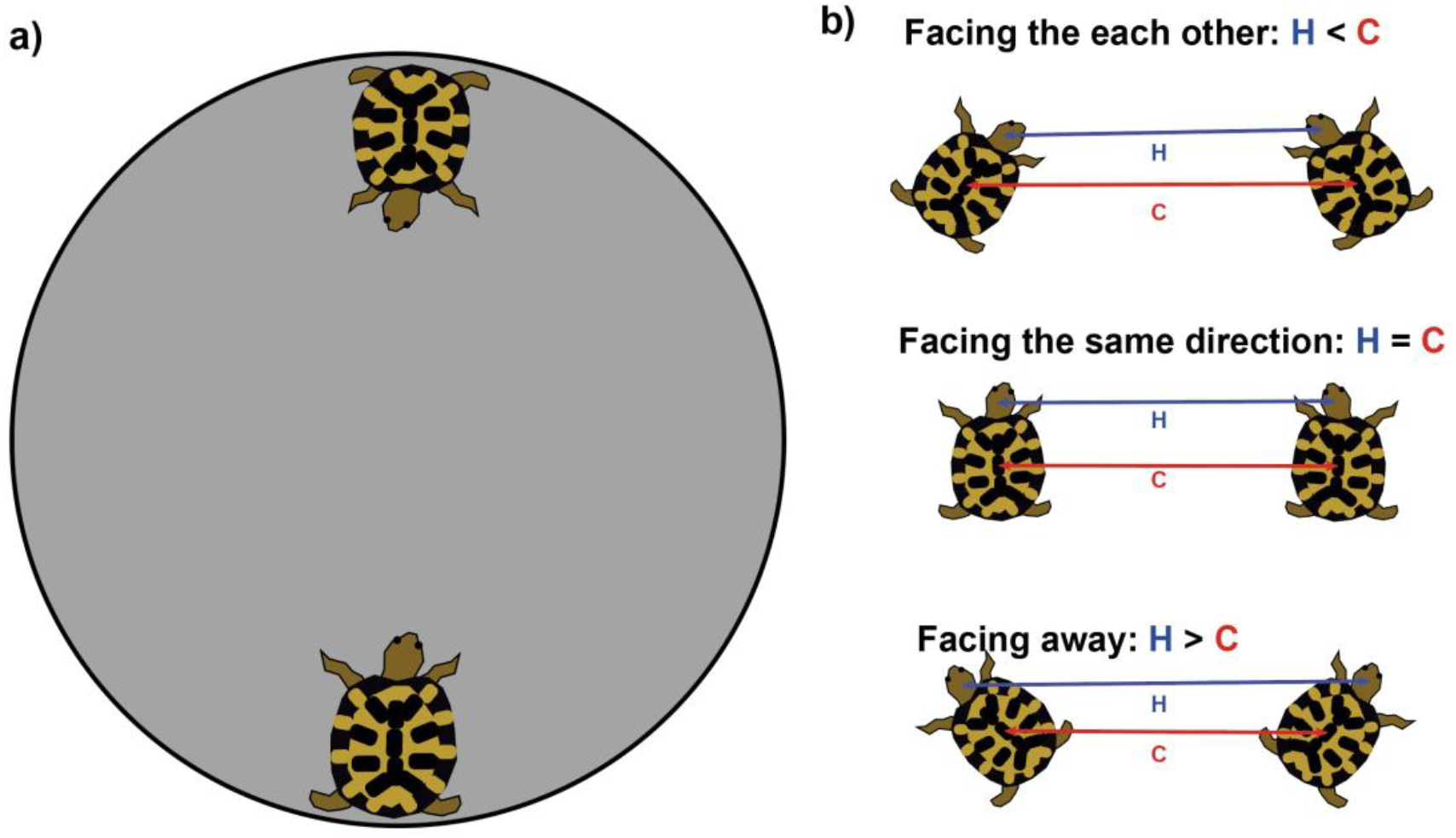
a) Schematic representation of the experimental apparatus as seen from above with subjects located in the starting position. b) Pairwise distance between heads (H) and carapaces (C) of the two subjects. When H<C, tortoises face each other; when H=C, tortoises are parallel, facing the same direction; when H>C, tortoises face away from each other.

### Expected chance distance with random trajectories

We used the average plastron length (15.7 cm) as a proxy for the diameter of a tortoise. To calculate the chance pairwise distance between tortoise centroids we implemented a Fermi-like estimation method. Tortoises were simulated as circles with radius 7.85 cm (half of the plastron length). We simulated the random positions of 100,000 pairs of tortoises uniformly distributed within an arena of radius 25.5 cm (the same size used in the experiments), obtaining a chance carapace distance of 39.10 cm. All simulations were run in R (version 4.5.1).

### Procedure

Before starting each trial, each tortoise was placed in an open box in the sunlight to thermo-conform their body temperature, a procedure already used in previous experiments to make sure the tortoises were active (Versace et al. 2018, 2020; Damini et al. 2021). Body temperature was measured using a digital thermometer pointed at the centre of the carapace. The animals were tested once the carapace temperature reached 32°C, a temperature that ensures activity without imposing heat-stress (Chelazzi and Calzolai 1986; Vujović et al. 2023).

The experiment was conducted during the summer months (June–September) on sunny days between 09:00 and 16:00, when the tortoises were most active. Trials were not carried out on overcast or rainy days, as the tortoises were inactive under those conditions. In each trial, two subjects were placed in diametrically opposed positions (the furthest possible distance within the arena), facing the centre of the arena (see Fig. 1a).

Video recordings of trials lasted 10-20 minutes, depending on when the first instance of movement occurred in one tortoise, allowing for max 10 minutes of inactivity. Each trial was analysed for 10 minutes from the first instance of movement (defined as displacement of the carapace following movement of a limb) (Versace et al. 2018). We chose a short duration of 10 minutes to minimise stress due to confinement. In this timeframe, no harmful behaviour (e.g. persistent biting or failed retreat) was observed. Aggressive actions, such as blows directed to the other animal, bite displays, and attempts to overturn the other individual were noted throughout the trial. At least 24 hours elapsed between the two trials for each subject. Following each trial, animals were returned to their home enclosures.

### Video and data analysis

#### Position and orientation

Using ImageJ (Rasband 2017), we recorded the coordinates of each individual’s carapace centroid and the tip of the head in each frame, using a strategy similar to the one used in previous studies with tortoise hatchlings (Versace et al. 2018; Damini et al. 2021),

For each 2-minute time bin (five time-bins per trial), we calculated: the Euclidean distance (cm) between the centroid of the two carapaces (C) and between the tips of the two heads (H), and the difference between these measures (H-C) (see Figure 1b). The H-C distance indicated whether the two subjects were facing each other (H<C), facing the same direction (H=C), or facing away from each other (H>C). We also computed a Facing orientation coefficient as (C–H) / (C+H), to standardise facing behaviour across varying proximity. A positive value of the coefficient indicated that tortoises are facing each other, a negative value that they are facing away from each other.

#### Statistical analysis of social cohesion/avoidance

We investigated the effects of Familiarity (familiar vs. unfamiliar), Sex (female vs. male), Time (categorized into 2-minute blocks: 0 (starting time), 1-2, 3-4, 5-6, 7-8, and 9-10 min) and their interactions on pair proximity. We used a linear mixed-effects model (*lmer*), including Pair, Subject A and Subject B as random intercepts. Model assumptions were confirmed via visual inspection of diagnostic plots. Facing orientation was analysed with the same linear mixed-effects model. Some heavy-tailedness was considered appropriate due to the robustness of linear mixed-effects frameworks. We used Bonferroni-Holm corrections to control for multiple testing in post-hoc comparisons between sexes and time points, and with the chance expected distance. Significance was set at α=0.05.

#### Aggressive behavior

We scored the frequency of aggressive behaviours described in the literature (Auffenberg 1977; Doody et al. 2021): bite attempts (when one subject displayed a bite movement by opening and closing their mouth directed towards the conspecific), carapace blows (when one subject hits the other with its carapace) and attempts to overturn the other individual (when one subject used its own carapace to attempt to flip the conspecific over). We analysed the effect of Sex Familiarity, and their interaction (fixed effects) on the total aggressive interactions (total count of bites, carapace blows, overturning attempts) using a Negative Binomial Generalized Linear Model, to account for overdispersion in the count data (Dispersion ratio > 11) using glm.nb *(MASS* package). All analyses were conducted in R.

#### Ethics

The experimental procedures were approved by the Ethical Committee of the Fondazione Museo Civico Rovereto (Italy), Prot. Num. 0000322. Experiments comply with the current Italian and European Community laws for the ethical treatment of animals and are in line with the ASAB/ABS Guidelines for the Use of Animals in Research.

## Results

### Pair proximity (distance between carapace centroids)

We found a significant effect of Time on carapace proximity distance (all t values for time intervals <−5.27, p<0.001), indicating a decrease of distance. A significant interaction was observed between Sex and Time (e.g., t=−3.34, p=0.001 for the 1–2 min interval; t=−3.51, p<0.001 for the 9–10 min interval). No significant effects were found for Familiarity (t=−0.47, p=0.644), nor for interaction between Familiarity, Sex, and Time (all p>0.05). Post-hoc comparisons showed significant sex differences in time intervals 1-2 and 9-10 (both p<0.01), see Table 2. During the test, male tortoises approached each other early in the trial and then maintained a relatively stable proximity (signficantly closer than expected by chance) throughout the test. In contrast, females approached each other slowly during the early time bins, and then moved apart, toward the end of the test, reaching the random proximity distance (see Fig. 2 and Table 3 for the results of Bonferroni-Holm test against chance distance).

**Table 2.** List of subjects, sex, and length of the plastron from the central part of the gular scutes, to the central part of the anal scutes.

| **Subject** | **Sex** | **Length (cm)** |
| --- | --- | --- |
| **1** | Male | 17.8 |
| **2** | Male | 17.5 |
| **3** | Male | 17 |
| **4** | Male | 18 |
| **5** | Male | 17.5 |
| **6** | Male | 15 |
| **7** | Male | 16.5 |
| **8** | Male | 14 |
| **9** | Male | 14.5 |
| **11** | Male | 12 |
| **12** | Male | 15 |
| **18** | Male | 13 |
| **25** | Female | 14.5 |
| **26** | Female | 14.5 |
| **30** | Female | 15 |
| **32** | Female | 17.5 |
| **33** | Female | 17 |
| **34** | Female | 17 |
| **35** | Female | 14.5 |
| **36** | Female | 16.5 |

**Table 3.** Post-hoc comparisons between sexes for the proximity results. Bold indicates significant results.

| Time | t | df | p-value | Holm p-value |
| --- | --- | --- | --- | --- |
| Time 0 | 0.480 | 10.204 | 0.641 |  |
| **Time 1-2** | **4.727** | **8.640** | **0.001** | **0.007** |
| Time 3-4 | 1.734 | 14.048 | 0.105 | 0.419 |
| Time 5-6 | 0.393 | 15.059 | 0.700 | 1.000 |
| Time 7-8 | 1.01910.852 | 0.332 | 0.990 |  |
| **Time 9-10** | **4.028** | **8.471** | **0.003** | **0.017** |

**Figure 2.**
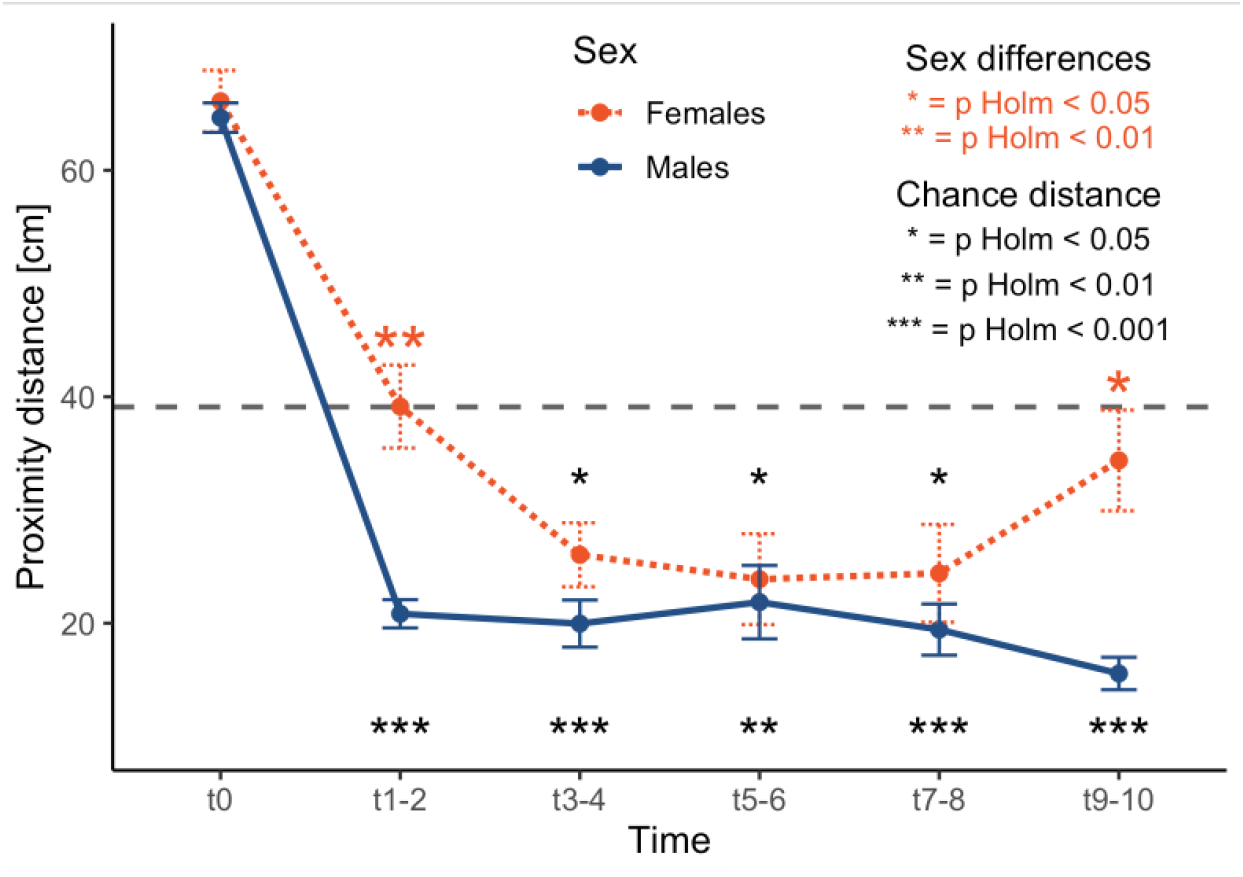
Average inter-individual distance (in cm, y axis) across the five time bins (1–2, 3–4, 5–6, 7– 8, 9–10 minutes, x axis), separated by sex. Blue lines represent males; red lines represent females. The dashed line represents the random expected distance. Subjects were initially placed in diametrically opposed locations (Time 0). Asterisks indicate significant differences between sexes (red colour) and against expected random expected distance for each sex at each time bin.

### Facing orientation

We found a significant main effect of Time on facing orientation (all t<−9.75, p<0.001), with a decrease in orientation towards each other compared to the initial baseline. A significant main effect of Sex was observed (t=2.46, p=0.016), with males orienting towards each other more than females, overall (see Figure 3). A significant Sex × Time interaction was also observed, specifically at the 3–4 min (t=−2.01, p=0.048), 7–8 min (t=−3.11, p=0.002), and 9–10 min (t=−2.04, p=0.044), with only the 7-8 minute difference remaining present after Bonferroni-Holm correction. No significant main effect was found for the Familiarity condition (t=1.50, p=0.136), nor for the three-way interaction between Familiarity, Sex, and Time (all p>0.05).

**Figure 3.**
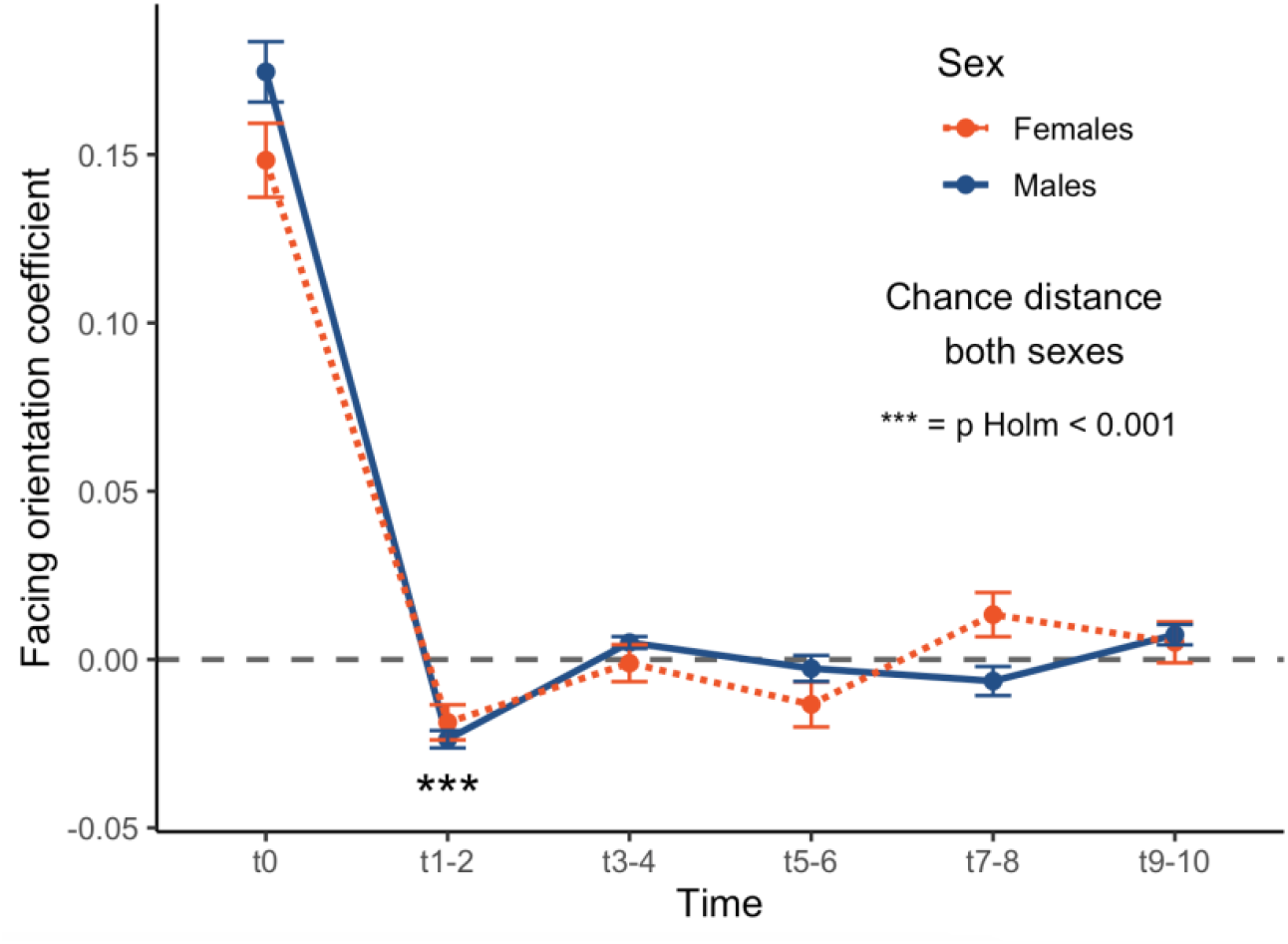
Average facing orientation coefficient (y axis) across five time bins (x axis), divided by sex. Positive values indicate that animals were facing each other; negative values indicate that they were facing away from each other. Blue lines represent males; red lines represent females. The dashed line represents the random expected orientation coefficient. Subjects were initially placed in diametrically opposed locations (Time 0). Asterisks indicate significant differences of the overall sample (females and males).

### Aggressive displays

Overall, across the 20 trials (12 male, 8 female pairs), we observed a total of 88 aggressive displays (see Table 4): 81 in males (mean=6.75/trial, mode=18), and 7 in females (mean=0.88/trial, mode=3). Sex had a significant positive effect on aggression (z=2.56, p=0.0104). Males displayed an eight-fold increase in aggressive interactions compared to females. Familiarity did not reach significance (z=−0.31, p=0.76). The high dispersion parameter (θ=0.415) reflects the uneven distribution of aggressive events among subjects, suggesting that a subset of individuals drove the majority of observed aggressive interactions.

**Table 4.** Post-hoc comparisons against chance expected distance by Sex and Time bin. Bold indicates significant results.

| Time/Sex | Average | t | df | p-value | Holm p-value |
| --- | --- | --- | --- | --- | --- |
| Time 0/Females | 66.105 | -9.914 | 7 | <0.001 |  |
| Time 0/Males | 64.655 | -19.69 | 11 | <0.001 |  |
| Time 1-2/Females | 39.140 | 0.543 | 7 | 0.604 | 0.650 |
| Time 1-2/Males | 20.832 | -14.642 | 11 | **<0.001** | **<0.001** |
| Time 3-4/Females | 26.039 | -4.629 | 77 | 0.002 | 0.012 |
| Time 3-4/Males | 9.964 | -9.194 | 11 | **<0.001** | **<0.001** |
| Time 5-6/Females | 23.887 | -3.794 | 7 | **0.007** | **0.027** |
| Time 5-6/Males | 21.860 | -5.314 | 11 | **<0.001** | **<0.001** |
| Time 7-8/Females | 24.407 | -3.400 | 7 | **0.011** | **0.034** |
| Time 7-8/Males | 19.436 | -8.683 | 11 | **<0.001** | **<0.001** |
| Time 9-10/Females | 34.379 | -1.059 | 7 | 0.325 | 0.649 |
| Time 9-10/Males | 15.558 | -16.420 | 11 | **<0.001** | **<0.001** |

**Table 5.** List of experimental pairs with sex, familiarity condition, and number of aggressive displays recorded during the test.

| Subj A | Subj B | Sex | Familiarity condition | Bite attempts | Carapace blows | Overturning attempts | Tot. |
| --- | --- | --- | --- | --- | --- | --- | --- |
| 33 | 34 | Female | Familiar | 0 | 0 | 0 | 0 |
| 35 | 25 | Female | Unfamiliar | 0 | 0 | 1 | 1 |
| 36 | 32 | Female | Unfamiliar | 1 | 0 | 0 | 1 |
| 26 | 30 | Female | Familiar | 2 | 1 | 0 | 3 |
| 30 | 34 | Female | Unfamiliar | 0 | 0 | 0 | 0 |
| 25 | 32 | Female | Familiar | 0 | 0 | 2 | 2 |
| 33 | 26 | Female | Unfamiliar | 0 | 0 | 0 | 0 |
| 35 | 36 | Female | Familiar | 0 | 0 | 0 | 0 |
| 4 | 5 | Male | Familiar | 4 | 0 | 0 | 4 |
| 6 | 12 | Male | Unfamiliar | 12 | 6 | 6 | 18 |
| 8 | 7 | Male | Familiar | 1 | 0 | 0 | 1 |
| 18 | 11 | Male | Unfamiliar | 5 | 0 | 0 | 5 |
| 1 | 3 | Male | Familiar | 1 | 0 | 0 | 1 |
| 9 | 7 | Male | Unfamiliar | 18 | 0 | 0 | 18 |
| 12 | 11 | Male | Familiar | 33 | 0 | 0 | 33 |
| 2 | 8 | Male | Unfamiliar | 0 | 0 | 0 | 0 |
| 6 | 18 | Male | Familiar | 0 | 0 | 0 | 0 |
| 1 | 4 | Male | Unfamiliar | 0 | 0 | 0 | 0 |
| 3 | 5 | Male | Unfamiliar | 1 | 0 | 0 | 1 |
| 2 | 9 | Male | Familiar | 0 | 0 | 0 | 0 |
| Overall |  | Female | Familiar | 2 | 1 | 2 | 5 |
| Overall |  | Female | Unfamiliar | 1 | 0 | 1 | 2 |
| Overall |  | Male | Familiar | 39 | 0 | 0 | 39 |
| Overall |  | Male | Unfamiliar | 36 | 6 | 12 | 42 |

## Discussion

We investigated whether adult captive tortoises (*Testudo hermanni*) behave differently when interacting with familiar versus unfamiliar same-sex conspecifics and how sex modules social strategies. After a two-month familiarisation phase with three conspecifics, tortoises were tested in a new arena with a familiar or with an unfamiliar conspecific. Contrary to our hypothesis, familiarity did not significantly influence inter-individual distance, orientation, or aggressive behaviours. In contrast, sex had a strong and significant impact in behavioural differences, independently from familiarity.

Male tortoises approached conspecifics rapidly and remained in close proximity throughout the trial, while females initially explored the other individual but progressively moved away, reaching the expected chance distance. This pattern mirrors what previously observed in unfamiliar tortoise hatchlings of closely related species (Versace et al. 2018), where hatchlings moved apart from unfamiliar animals each other after an initial exploration. However, differently from that study, our adult *Testudo hermanni* tortoises did not ignore familiar individuals.

Males and females differed also in facing orientation, with males facing each other significantly more than females, suggesting again less engagement of females in social interactions. The data on aggression, where males engaged significantly more than females in aggressive displays, suggests that the approach responses in males were likely motived by dominance and/or territorial goals. Overall, sex differences indicate that, at least in captive contexts, adult male and female tortoises are differently motivated in same-sex interactions, with males displaying strong exploratory and possibly competitive tendencies, and females a moderate initial interest, followed by disengagement. It’s interesting to notice that, in similar tests, social disengagement is typically not observed in social species. For instance, domestic chicks are attracted to conspecifics from the beginning of life, with a strong reinstatement motivation (the motivation to reach the group and remain in contact with conspecifics) (Versace et al. 2017, 2021; Pallante et al. 2021). In a similar test, chicks from different breeds quickly approach conspecifics and then remained closer to familiar compared to unfamiliar conspecifics (Versace et al. 2021).

We designed a short test, to minimise stress and avoid risks and interruptions caused by potential aggression. During the 10-minute trials, we did not observe behaviours that could have harmed the tortoises. Nevertheless, the presence of some aggressive displays, particularly among males, and the progressive reduction of distance between males, that remained significantly closed than expected by chance throughout the test, suggest that a longer test might have increased the risk of escalation. In this sense, the duration of 10 minutes appears appropriate for preventing potentially harmful interactions. More broadly, these observations highlight the importance of considering space requirements for tortoises in relation to sex and individual behavioural traits. We found that males showed significantly more aggressive displays than females, regardless of familiarity. Each tortoise was observed only twice: once with a familiar individual and once with an unfamiliar one, making it difficult to draw firm conclusions about consistent individual behavioural tendencies. Nonetheless, aggressive displays were not evenly distributed across individuals. A small subset of tortoises accounted for most interactions (e.g., Subject 12 was particularly proactive in initiating aggressive displays). Although these patterns may indicate that male tortoises experience greater stress than females in confined spaces, and that individual differences are relevant, the limited sample size of our study requires caution in interpretation. Future research with short trials and larger samples should further investigate the presence of potential aggressive displays to clarify sex differences and better understand how tortoises use these interactions to establish dominance or make territorial claims. Such information could help inform space allocation strategies in zoos and bio-parks, where higher housing densities have been associated with increased aggressive displays (O’Brien et al. 2025).

The lack of differential behaviour toward familiar and unfamiliar conspecifics surprised us. This result is compatible with several non-exclusive possibilities. It is possible that in a novel environment, adult tortoises are primarily driven by territorial motivations, which may override recognition-based social strategies such as the “dear enemy effect” (Ydenberg et al. 1988). In this novel context, males may attempt to assert dominance, while females may seek to increase spatial dispersal, avoiding potential resource competition or conflict. Such a difference in motivation between males and females is supported by the similarity of responses between females and tortoise hatchlings previously tested in a similar task with unfamiliar individuals (Versace et al. 2018), where we observed approach followed by avoidance. That study, however, was performed on *Testudo graeca* and *Testudo marginata* animals. As we are not aware of other studies testing familiarity recognition in *T. hermanni*, it is possible that our study subjects were not able to differentiate between familiar and unfamiliar conspecifics. We consider this scenario unlikely. Given that *Testudo* hatchlings as young as four weeks recognise familiar individuals (Versace et al. 2018), it appears unlikely that adult *T. hermanni* don’t possess this ability. The differences observed between closely related *Testudo* hatchlings (*T. graeca, T. marginata*) and adult behaviour observed here in *T. hermanni* may reflect developmental shifts in ecological priorities, such as mating competition or territory defence. Another possibility is that familiarity recognition skills are present but not expressed in these conditions. This would parallel observations conducted in a study published during our paper revision (Goessling et al. 2026), where headstart Gopher tortoises (*Gopherus polyphemus*) did not modulate their social behaviour (nipping, chasing, headbobbing, and colliding) based on familiarity but responded differently to familiar/unfamiliar odourants. Another possibility is that, differently from the less experienced *Testudo* hatchlings, these adult tortoises were confused by subtle olfactory cues, as all pairs were tested in the same room and Hermann’s tortoises are able to use olfaction in social contexts (Galeotti et al. 2007). It is also plausible that tortoises’ behaviour is more strongly modulated by sex than familiarity, especially for a captive population used to encounter other individuals on a daily basis, and that the extended captive history of our subjects (at least three years, typically longer) and prior housing with other tortoises, may have influenced these behavioural choices. Future studies conducted on animals that have not spent long time in captivity should clarify whether there are any habituation or sensitisation effects induced by captivity. Sex-specific strategies in adults may hence mask familiarity effects. Moreover, since our observation window was limited to 10 minutes, to minimise stress in a restricted location, this setting may have limited the expression of subtler social dynamics, such as escalation or resolution of conflict via avoidance. Future studies should address these points.

Taken together, our findings reinforce the view that tortoises are able to actively pursue different social strategies, from exploration of other individuals to continuous engagement and maintaining moving away from other individuals. However, while *Testudo* hatchlings differentiate familiar from unfamiliar conspecifics and actively avoided the latter (Versace et al., 2018), adults did not appear to do so in our setting, suggesting that ecological factors such as sexual maturity, territoriality, or context-specific behavioural regulation may influence the expression of social cohesion and social interactions in adulthood.

The sex differences observed in this study echo prior research on physiological and behavioural traits in tortoises. For instance, sex differences in corticosterone levels (Lance et al. 2001), patterns of burrow use (Bulova 1994), olfactory perception (Galeotti et al. 2007) have been reported. These sexual differences, in some cases, have also informed conservation efforts. It has been shown that male tortoises tend to disperse further following translocation, while females exhibit higher mortality rates than males (Germano et al. 2017). Our study shows that captive tortoises differ in same-sex interactions, and that males might be exposed to more stressful social interactions in confined locations. Recognising such sex-specific behaviours may therefore be critical for improving conservation planning and post-release monitoring.

About half of chelonians species are considered “Threatened” by the IUCN List criteria (Rhodin et al. 2018; Chen et al. 2025). Further studies of the social behaviour of wild and captive tortoises, and in particular how sex differences influence their interaction with conspecifics outside of the mating context, could improve our understanding of these animals and inform conservation efforts (Doody, 2023) in the vulnerable species *T. hermanni* studied here, and in other species.

In conclusion, while we did not find evidence of adult tortoises modulating their behaviour based on familiarity, we observed that sex strongly shapes patterns of spatial interaction and aggression in same-sex encounters. These behavioural differences highlight how sex should be taken into account in future studies, tortoise housing and conservation planning.

## Authors’ contributions

E.V., G.S., and S.D. conceived and designed the experiment; S.D. carried out the experiment; S.D. analysed the data and curated the dataset; S.D. and E.V. drafted the paper; all authors revised the manuscript and gave final approval for publication.

## Acknowledgments

We thank the internship students from the high school “Liceo Antonio Rosmini” (Rovereto, Italy) for help in data collection, and the Rovereto Civic Museum Foundation, for providing the facilities and the support to carry out this research.

## References

Aragón P, López P, Martín J (2003) Differential Avoidance Responses to Chemical Cues from Familiar and Unfamiliar Conspecifics by Male Iberian Rock Lizards (Lacerta monticola). Journal of Herpetology 37:583–585. 10.1670/192-02n

Auffenberg W (1977) Display behavior in tortoises. American Zoologist 17:241–250. 10.1093/icb/17.1.241

Bee MA, Gerhardt HC (2002) Individual voice recognition in a territorial frog (Rana catesbeiana). Proc R Soc Lond B 269:1443–1448. 10.1098/rspb.2002.2041

Bulova SJ (1994) Patterns of Burrow Use by Desert Tortoises: Gender Differences and Seasonal Trends. Herpetological Monographs 8:133. 10.2307/1467077

Burghardt GM (2013) Environmental enrichment and cognitive complexity in reptiles and amphibians: Concepts, review, and implications for captive populations. Applied Animal Behaviour Science 147:286–298. 10.1016/j.applanim.2013.04.013

Chelazzi G, Calzolai R (1986) Thermal benefits from familiarity with the environment in a reptile. Oecologia 68:557–558. 10.1007/BF00378771

Chen C, Wang J, Holyoak M, et al (2025) Global assessment of current extinction risks and future challenges for turtles and tortoises. Nat Commun 16:7114. 10.1038/s41467-025-62441-2

Cheney DL, Seyfarth RM (1980) Vocal recognition in free-ranging vervet monkeys. Animal Behaviour 28:362–367. 10.1016/S0003-3472(80)80044-3

Damini S, Stancher G, Versace E (2021) Recognition of familiar objects in tortoise hatchlings (Testudo spp.)

Doody JS, Burghardt GM, Dinets V (2013) Breaking the Social–Non-social Dichotomy: A Role for Reptiles in Vertebrate Social Behavior Research? Ethology 119:95–103. 10.1111/eth.12047

Doody JS, Dinets V, Burghardt GM (2021) The secret social lives of reptiles. Johns Hopkins University Press, Baltimore

Ebensperger LA, Cofré H (2001) On the evolution of group-living in the New World cursorial hystricognath rodents. Behavioral Ecology 12:227–236. 10.1093/beheco/12.2.227

Ernst CH, Barbour RW (1989) Turtles of the world. Smithsonian Institution Press, Washington DC

Freeland L, Ellis C, Michaels CJ (2020) Documenting Aggression, Dominance and the Impacts of Visitor Interaction on Galápagos Tortoises (Chelonoidis nigra) in a Zoo Setting. Animals 10:699. 10.3390/ani10040699

Galeotti P, Sacchi R, Pellitteri-Rosa D, Fasola M (2005) Female preference for fast-rate, high-pitched calls in Hermann’s tortoises Testudo hermanni. Behavioral Ecology 16:301–308. 10.1093/beheco/arh165

Galeotti P, Sacchi R, Rosa DP, Fasola M (2007) Olfactory Discrimination of Species, Sex, and Sexual Maturity by the Hermann’s Tortoise Testudo Hermanni. Copeia 2007:980–985. 10.1643/0045-8511(2007)7%255B980:ODOSSA%255D2.0.CO;2

Germano JM, Nafus MG, Perry JA, et al (2017) Predicting translocation outcomes with personality for desert tortoises. Behavioral Ecology 28:1075–1084. 10.1093/beheco/arx064

Goessling JM, Weikert G, Conrad L, Hilton ML (2026) Old neighbours, long-lost siblings, and total strangers: Social environment impacts on headstart tortoise behaviours. Applied Animal Behaviour Science 294:106869. 10.1016/j.applanim.2025.106869

Hofmann H a., Beery AK, Blumstein DT, et al (2014) An evolutionary framework for studying mechanisms of social behavior. Trends in Ecology & Evolution 1–9. 10.1016/j.tree.2014.07.008

Husak JF, Fox SF (2003) Adult male collared lizards, Crotaphytus collaris, increase aggression towards displaced neighbours. Animal Behaviour 65:391–396. 10.1006/anbe.2003.2058

IUCN (2024) Testudo hermanni: Luiselli, L.: The IUCN Red List of Threatened Species 2024: e.T21648A268511652

Kendrick KM, da Costa AP, Leigh AE, et al (2001) Sheep don’t forget a face. Nature 414:165–166. 10.1038/35102669

Klaphake E (2010) A Fresh Look at Metabolic Bone Diseases in Reptiles and Amphibians. Veterinary Clinics of North America: Exotic Animal Practice 13:375–392. 10.1016/j.cvex.2010.05.007

Krause J, Ruxton GD (2002) Living in Groups. Oxford University Press

Lance VA, Grumbles JS, Rostal DC (2001) Sex differences in plasma corticosterone in desert tortoises,Gopherus agassizii, during the reproductive cycle. J Exp Zool 289:285–289. 10.1002/1097-010x(20010415/30)289:5%253C285::aid-jez2%253E3.0.co;2-b

Madile Hjelt M, Moyano L, Echave ME, et al (2024) Social networks of threatened Chaco tortoises (Chelonoidis chilensis) in the wild. Biological Journal of the Linnean Society 143:blae073. 10.1093/biolinnean/blae073

Makuya L, Schradin C (2024) The secret social life of solitary mammals. Proc Natl Acad Sci USA 121:. 10.1073/pnas.2402871121

Martín J, Raya-García E, Ortega J, López P (2021) Offspring and adult chemosensory recognition by an amphisbaenian reptile may allow maintaining familiar links in the fossorial environment. PeerJ 9:e10780. 10.7717/peerj.10780

Mazzotti S, Fasola M, Pisapia A (2002) Activity and home range of Testudo hermanni in Northern Italy. Amphibia-Reptilia 23:305–312. 10.1163/15685380260449180

Munch KL, Noble DWA, Wapstra E, While GM (2018) Mate familiarity and social learning in a monogamous lizard. Oecologia 188:1–10. 10.1007/s00442-018-4153-z

O’Brien SL, Diaz A, Cronin KA (2025) Social dynamics and behavior of zoo-housed red-footed tortoises at different housing densities. Behavioural Processes 231:105242. 10.1016/j.beproc.2025.105242

Pallante V, Rucco D, Versace E (2021) Young chicks quickly lose their spontaneous preference to aggregate with females. Behav Ecol Sociobiol 75:78. 10.1007/s00265-021-03012-5

Pearse DE, Avise JC (2001) Turtle Mating Systems: Behavior, Sperm Storage, and Genetic Paternity. Journal of Heredity 92:206–211. 10.1093/jhered/92.2.206

Rasband W (2017) ImageJ. U S National Institutes of Health, Bethesda, Maryland, USA //imagej.nih.gov/ij/

Ray EJ, Maruska KP (2023) Sensory Mechanisms of Parent-Offspring Recognition in Fishes, Amphibians, and Reptiles. Integrative And Comparative Biology 63:1168– 1181. 10.1093/icb/icad104

Rhodin AGJ, Stanford CB, Dijk PPV, et al (2018) Global Conservation Status of Turtles and Tortoises (Order Testudines). Chelonian Conservation and Biology 17:135. 10.2744/ccb-1348.1

Rubenstein DR, Abbot P (eds) (2017) Comparative Social Evolution. Cambridge University Press, Cambridge

Sacchi R, Galeotti P, Fasola M, Ballasina D (2003) Vocalizations and courtship intensity correlate with mounting success in marginated tortoises Testudo marginata. Behavioral Ecology and Sociobiology 55:95–102. 10.1007/s00265-003-0685-1

Sah P, Nussear KE, Esque TC, et al (2016) Inferring social structure and its drivers from refuge use in the desert tortoise, a relatively solitary species. Behav Ecol Sociobiol 70:1277–1289. 10.1007/s00265-016-2136-9

Salguero-Gómez R (2024) More social species live longer, have longer generation times and longer reproductive windows. Phil Trans R Soc B 379:. 10.1098/rstb.2022.0459

Santacà M, Wilkinson A, Stancher G, et al (2025) Lizards and tortoises show evidence of low inhibitory control. Sci Rep 15:23446. 10.1038/s41598-025-08373-9

Semaha MJ, Rodríguez-Caro RC, Giménez A, et al (2025) Captive-introduced tortoises in wild populations: can we identify them by shell morphology? Eur J Wildl Res 71:13. 10.1007/s10344-024-01893-1

Sheehan MJ, Tibbetts EA (2011) Specialized face learning is associated with individual recognition in paper wasps. Science 334:1272–1275. 10.1126/science.1211334

Silk JB (2007) The adaptive value of sociality in mammalian groups. Philosophical Transactions of the Royal Society B: Biological Sciences 362:539–559. 10.1098/rstb.2006.1994

Stubbs D, Swingland IR (1985) The ecology of a Mediterranean tortoise (Testudo hermanni): a declining population. Can J Zool 63:169–180. 10.1139/z85-026

Tibbetts EA, Dale J (2007) Individual recognition: it is good to be different. Trends in Ecology and Evolution 22:529–537. 10.1016/j.tree.2007.09.001

Twining JP, Sutherland C, Zalewski A, et al (2024) Using global remote camera data of a solitary species complex to evaluate the drivers of group formation. Proc Natl Acad Sci USA 121:. 10.1073/pnas.2312252121

Versace E, Damini S, Caffini M, Stancher G (2018) Born to be asocial: newly hatched tortoises avoid unfamiliar individuals. Animal Behaviour 138:187–192. 10.1016/j.anbehav.2018.02.012

Versace E, Damini S, Stancher G (2020) Early preference for face-like stimuli in solitary species as revealed by tortoise hatchlings. Proceedings of the National Academy of Sciences of the United States of America 117:. 10.1073/pnas.2011453117

Versace E, Fracasso I, Baldan G, et al (2017) Newborn chicks show inherited variability in early social predispositions for hen-like stimuli. Scientific Reports 7:40296. 10.1038/srep40296

Versace E, Ragusa M, Pallante V, Wang S (2021) Attraction for familiar conspecifics in young chicks (Gallus gallus): An interbreed study. Behavioural Processes 193:104498. 10.1016/j.beproc.2021.104498

Vujović A, Pešić V, Meek R (2023) Living in a Thermally Diverse Environment: Field Body Temperatures and Thermoregulation in Hermann’s Tortoise, Testudo hermanni, in Montenegro. Conservation 3:59–70. 10.3390/conservation3010005

Ward A, Webster M (2016) Sociality: The Behaviour of Group-Living Animals. Springer International Publishing, Cham

Ward AJW, Kent MIA, Webster MM (2020) Social Recognition and Social Attraction in Group-Living Fishes. Front Ecol Evol 8:. 10.3389/fevo.2020.00015

Wendland LD, Wooding J, White CL, et al (2010) Social behavior drives the dynamics of respiratory disease in threatened tortoises. Ecology 91:1257–1262. 10.1890/09-1414.1

Wilkinson A, Kuenstner K, Mueller J, Huber L (2010a) Social learning in a non-social reptile (Geochelone carbonaria). Biology letters 6:614–616. 10.1098/rsbl.2010.0092

Wilkinson A, Mandl I, Bugnyar T, Huber L (2010b) Gaze following in the red-footed tortoise (Geochelone carbonaria). Animal Cognition 13:765–9. 10.1007/s10071-010-0320-2

Ydenberg RC, Giraldeau LA, Falls JB (1988) Neighbours, strangers, and the asymmetric war of attrition. Animal Behaviour 36:343–347. 10.1016/S0003-3472(88)80004-6

